# Projection-specific visual feature encoding by layer 5 cortical subnetworks

**DOI:** 10.1101/028910

**Authors:** Gyorgy Lur, Martin A. Vinck, Lan Tang, Jessica A. Cardin, Michael J. Higley

**Affiliations:** Department of Neuroscience; Program in Cellular Neuroscience, Neurodegeneration and Repair

## Abstract

Primary neocortical sensory areas act as central hubs, distributing afferent information to numerous cortical and subcortical structures. However, it remains unclear whether each downstream target receives distinct versions of sensory information. We used *in vivo* calcium imaging combined with retrograde tracing to monitor visual response properties of three distinct subpopulations of projection neurons in primary visual cortex. While there is overlap across the groups, on average corticotectal (CT) cells exhibit lower contrast thresholds and broader tuning for orientation and spatial frequency in comparison to corticostriatal (CS) cells, while corticocortical (CC) cells have intermediate properties. Noise correlational analyses support the hypothesis that CT cells integrate information across diverse layer 5 populations, whereas CS and CC cells form more selectively interconnected groups. Overall, our findings demonstrate the existence of functional subnetworks within layer 5 that may differentially route visual information to behaviorally relevant downstream targets.

## Introduction

Recent evidence suggests that transmission of sensory information over distinct channels to different downstream targets is a key feature of cortical circuits (Wang and Burkhalter, 2013). Indeed, primary sensory cortex may act as a hub for routing information streams from a locally heterogeneous population of pyramidal neurons (PNs) (Glickfeld et al., 2013, Jarosiewicz et al., 2012). However, the extent to which pools of PNs extract distinct feature information from sensory inputs remains unclear. The relationships between sensory processing and functional connectivity within local and long distance cortical networks are also poorly understood.

In the visual cortex, connection probability is elevated for neurons sharing similar feature selectivity (Ko et al., 2011, Kohn and Smith, 2005, Okun et al., 2015). However, this relationship between connectivity and sensory tuning is not exclusive, as not all connected neurons respond to identical features (Ko et al., 2014). In addition, not all connected neurons share the same target structures (Brown and Hestrin, 2009). Along with diverse intracortical projections, V1 projects heavily from layers 2/3 and 5 to subcortical structures, including the basal ganglia and tectum (Khibnik et al., 2014, Oh et al., 2014).

Data from *ex vivo* preparations suggests that different populations of PNs in layer 5 (L5) may be functionally distinct. For example, corticotectal (CT) neurons projecting to the superior colliculus have thick apical trunks with prominent dendritic tuft arborizations and express high levels of HCN channels (Harris and Shepherd, 2015, Kasper et al., 1994). In contrast, non-CT cells, including corticostriatal (CS) and corticocortical (CC) neurons, have more modest apical dendritic tufts and exhibit little HCN channel expression (Shepherd, 2013, Larkman and Mason, 1990). Moreover, distinct L5 populations are differentially connected with superficial layers and with each other, suggesting the existence of distinct subnetworks within neocortical circuits (Lefort et al., 2009, Feldmeyer, 2012). Indeed, in mouse visual cortex, intra-group synaptic connectivity is highest for CS cells, contrasting with CT cells that broadly receive inputs from diverse L5 populations (Brown and Hestrin, 2009).

Previous *in vivo* work has shown that, in general, L5 neurons are more broadly tuned for orientation and spatial frequency than neurons in more superficial layers (Niell and Stryker, 2008, Hoy and Niell, 2015). However, it is less clear how visual response properties vary across distinct cellular populations in L5. The striatum and superior colliculus are postulated to play important yet distinct roles in visually guided behavior (Sahibzada et al., 1986, Ragozzino et al., 2002), and the nature of the visual information directed to these areas from V1 is unclear. One possibility is that subcortical structures all receive a composite visual output, maximizing the efficacy and redundancy of visual signal transmission. Alternatively, subcortical projections may provide target-specific information content about visual features in the environment.

To address this issue, we combined retrograde fluorescent labeling with *in vivo* multiphoton calcium imaging to compare visual feature extraction across identified L5 PN populations. We find that CS, CC, and CT cells comprise largely non-overlapping populations in L5 of mouse V1. Furthermore, CT cells are more sensitive to low contrast and are more broadly tuned for orientation and spatial frequency than CS cells, while CC cells exhibit intermediate properties. Both CS and CC cells exhibit strong intra-group correlational structure, suggesting they form distinct subnetworks in L5, whereas CT cells show broad correlations across groups. These findings indicate that visual features may be differentially extracted by target-specific subnetworks of L5 PNs that route behaviorally relevant information to divergent downstream areas.

## Results

### Distinct populations of pyramidal neurons in V1 layer 5

Previous studies have suggested that layer 5 comprises diverse groups of pyramidal neurons (PNs) that differ in their projection targets, morphology, and electrophysiological characteristics (Hattox and Nelson, 2007, Shepherd, 2013, Harris and Shepherd, 2015, Kasper et al., 1994, Larkman and Mason, 1990). To investigate the distinct functional properties of layer 5 PN subpopulations in V1, we combined fluorescent retrograde labeling with *in vivo* 2-photon calcium (Ca2+) imaging in lightly anesthetized mice (Fig. 1A-C, Fig. S1A,B). We identified three separate groups of PNs by injecting the retrograde tracer cholera toxin B (CTB) into either the superior colliculus (SC), dorsal striatum (dStr), or contralateral medial V2 (cV2) (Fig. S1C, see Methods). Using double injections of green and red fluorescent CTB, we confirmed that labeled populations in V1 are largely non-overlapping (<2% overlap) for the three classes (Fig. S1D-E), which also differed in their morphology and intrinsic electrophysiological characteristics (Fig. S1G-I and Table S1). Notably, corticotectal (CT), corticostriatal (CS), and corticocortical (CC) cells showed considerable overlap in their distribution as a function of cortical depth (Fig. S1F).

### Visual feature encoding by layer 5 PNs

For functional imaging, we injected red fluorescent CTB into one of the three target areas and expressed GCaMP6s (Chen et al., 2013) in V1 using a viral vector. We imaged 1525 neurons in 20 animals, of which 1279 were deemed visually responsive (see Methods). Of these, 950 were identified by tracer injection (342 CT cells from 6 animals; 306 CC neurons from 9 animals; 302 CS cells from 5 animals). The fraction of visually responsive cells was similar in all 3 populations (CT: 83%, CC: 80% and CS: 83%). Each cell was imaged during presentation of one or more visual stimulus sequences, consisting of whole-field sinusoidal drifting gratings with varied contrast, orientation, and spatial frequency. Importantly, *ex vivo* imaging revealed no differences across cell types with regard to the relationship between spiking and calcium signal (Fig. S2A-C).

Consistent with previous recordings of both spiking and sub-threshold activity (Hubel and Wiesel, 1962, Mechler and Ringach, 2002, Skottun et al., 1991), we observed cells whose visually-evoked Ca2+ transients were modulated to differing degrees at the temporal frequency of the grating stimulus. We quantified this property using a modulation index (MI, see Methods). Cells with higher MI values are more simple-like, while those with lower values are more complex-like (Fig. 1D). Using this metric, CT cells showed significantly weaker modulation (0.448±0.018, n=115, 6 animals) in comparison to CC (0.56±0.051, n=116, 9 animals, p=0.02, Student’s t-test) and CS (0.582±0.037, n=104, 5 animals, p=0.0006, t-test; Fig. 1E, Fig. S3A). There was no difference between CC and CS cells (p=0.36, t-test). Moreover, the period of the best-fit sine wave for the data was 0.9±0.2s, in agreement with the 1 Hz temporal frequency of the stimulus. There was no significant correlation between Ca2+ decay and the MI (Pearson’s r=-0.045, p=0.3273; Fig. S2D), suggesting that disparate Ca2+ buffering did not contribute to the observed MI differences. Importantly, we also found no significant differences between the decay kinetics of the Ca2+ signal across populations, suggesting that GCaMP6 expression is similar in the different cell groups (Fig. S2E).

We then measured the sensitivity to stimulus contrast across cell populations. Only cells with a significant contrast-dependent increase in response magnitude were considered for analysis (272/438 cells, Spearman rank test r>0 and p<0.05, Figure 1F). For each cell, we fitted the data with a hyperbolic ratio function (Figure 1G, see Methods) (Contreras and Palmer, 2003). We calculated the c50 value, exponent, and Rmax for the resulting curves with goodness-of-fit R^2^ values >0.4. The c50 value of CT cells (34.44±3.4%, n=70 cells, 6 animals) was significantly lower than that of CS cells (43.85±2.3%, n=75 cells, 5 animals, p=0.0112, Student’s t-test) or CC cells (43.14±3 %, n=108 cells, 9 animals, p=0.0285, t-test; Fig. 1H, Fig. S3B). Again, there was no difference between CS and CC cells (p=0.426, t-test). The exponent value was significantly higher in CT neurons (6.96±0.8) than in CC cells (5.27±0.57, p=0.043, t-test) but was not statistically different from CS cells (5.36±0.65, p=0.060, t-test, Fig. 1I, Fig. S3C). There was no significant difference between CC and CS cells (p=0.45, t-test). On average, CC cells exhibited a higher Rmax value (0.536±0.042) then CT (0.431+0.027, p=0.017, t-test) or CS cells (0.411±0.06, p=0.043, t-test; Fig. S3D,E). Together, these data indicate that, on average, CT cells are more complex-like and have a lower threshold for detecting visual stimuli compared with CS or CC cells.

**Figure 1.**
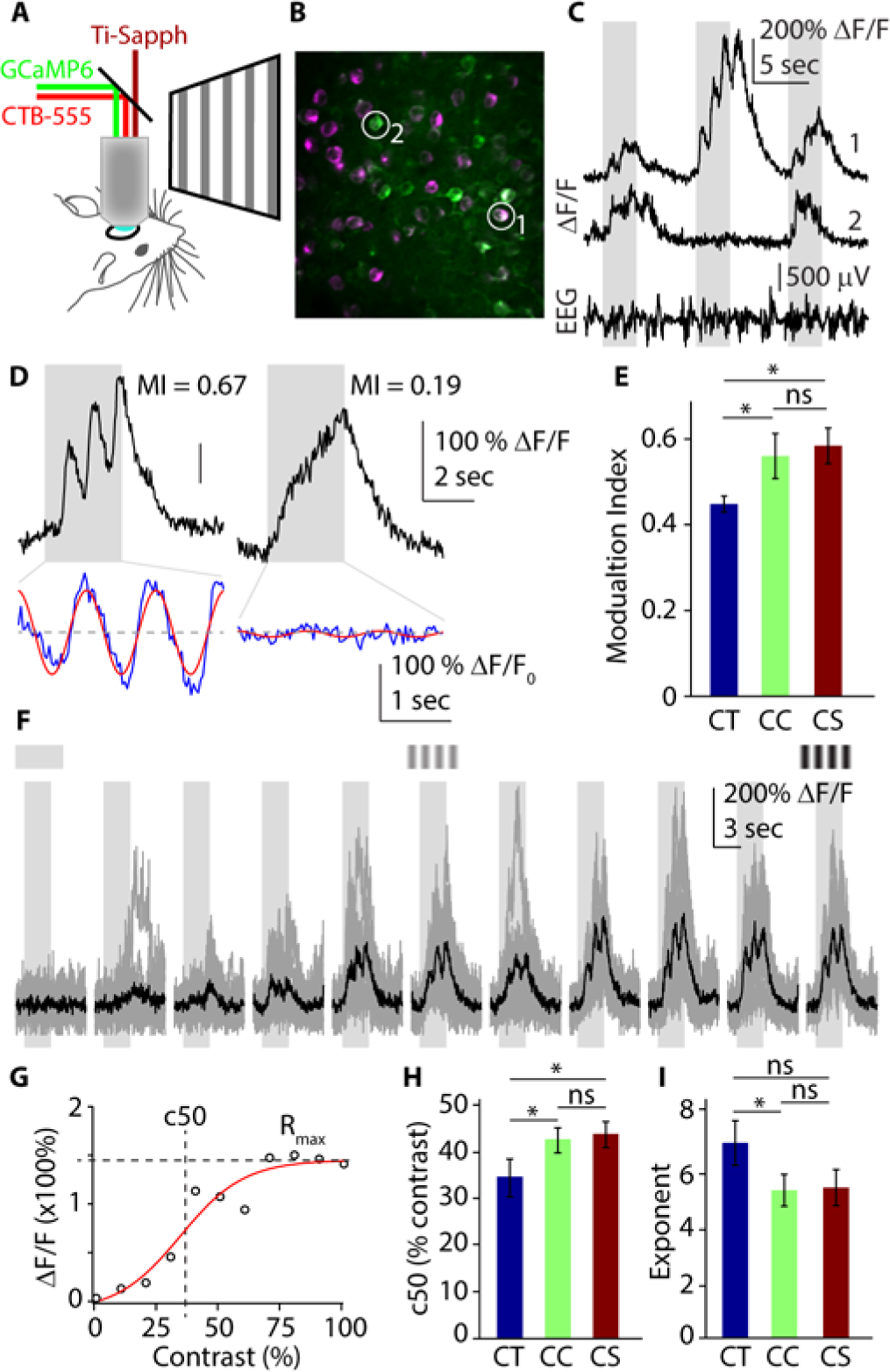
CT cells exhibit lower visual detection threshold than CC and CS neurons. (A) Schematic of *in vivo* 2-photon Ca2+ imaging of labeled L5 PN populations. (B) Example field of view. Green somata express GCaMP6s. Magenta cells express GCaMP6s and are retrogradely labeled with red fluorescent CTB-Alexa Fluor-555. (C) Example raw traces recorded from cells indicated in (B) and corresponding EEG signal. (D) Example ∆F/F traces (black) and de-trended visual responses (blue) with best fit sine waves (red) to calculate modulation index (MI). (E) Bars represent mean ± SEM modulation index for CT (blue) CC (green) and CS (red) cells. (F) Example raw (gray) and average (black) ∆F/F traces recorded at varying contrast values. (G) Hyperbolic ratio function fit (red) to contrast response (black circles). Dashed lines highlight c50 and R_max_ points. (H) Bars represent mean ± SEM c50 values of CT (blue) CC (green) and CS (red) cells. (I) Bars represent mean ± SEM exponent values of CT (blue) CC (green) and CS (red) cells. *: p<0.05, Student’s t-test, semi-weighted statistics (see Methods).

**Figure 2.**
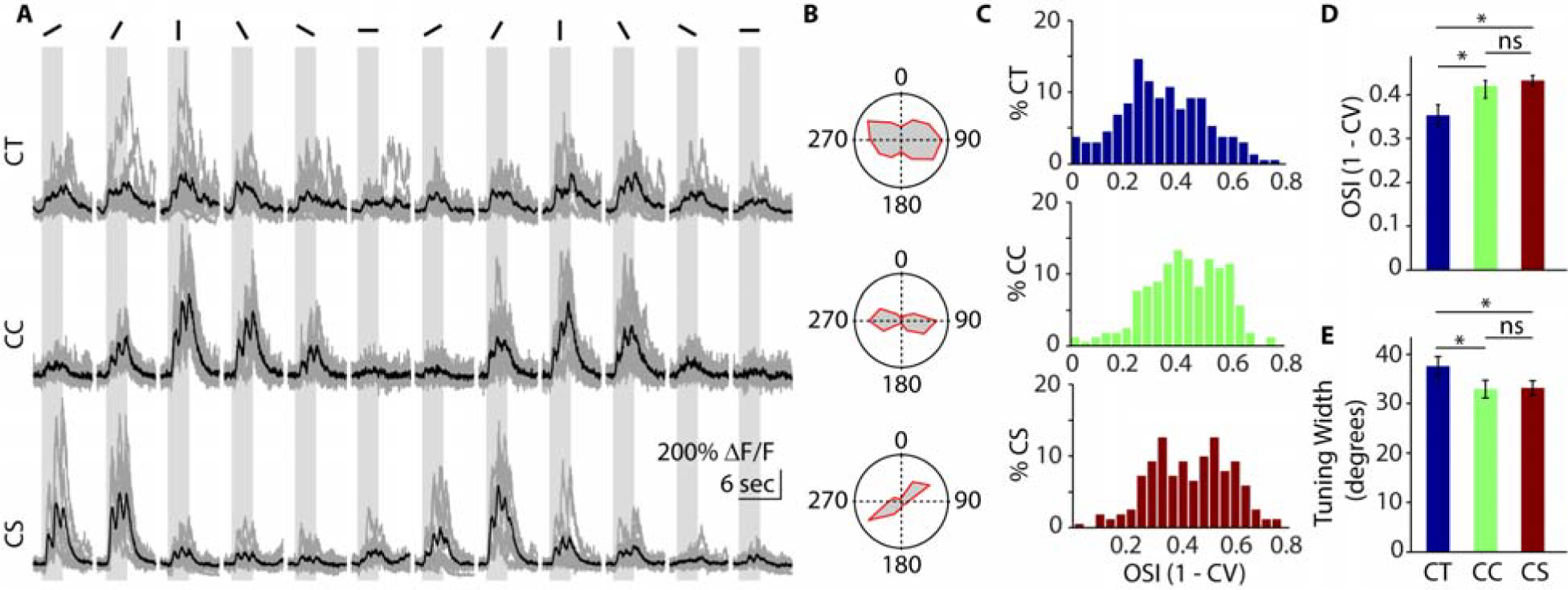
CT neurons are more broadly tuned for orientation than CC and CS cells. (A) Example raw (gray) and average (black) traces of CT (top), CC (middle) and CS (bottom) neurons at varying orientations. (B) Polar plots indicating the orientation tuning of the cells in (A). (C) Distribution of OSI values for CT, CC, and CS populations. (D) Bars represent mean ± SEM OSI values of CT (blue) CC (green) and CS (red) cells. (E) Bars represent mean ± SEM orientation tuning width of CT (blue) CC (green) and CS (red) cells. *: p<0.05, Student’s t-test, semi-weighted statistics (see Methods).

**Figure 3.**
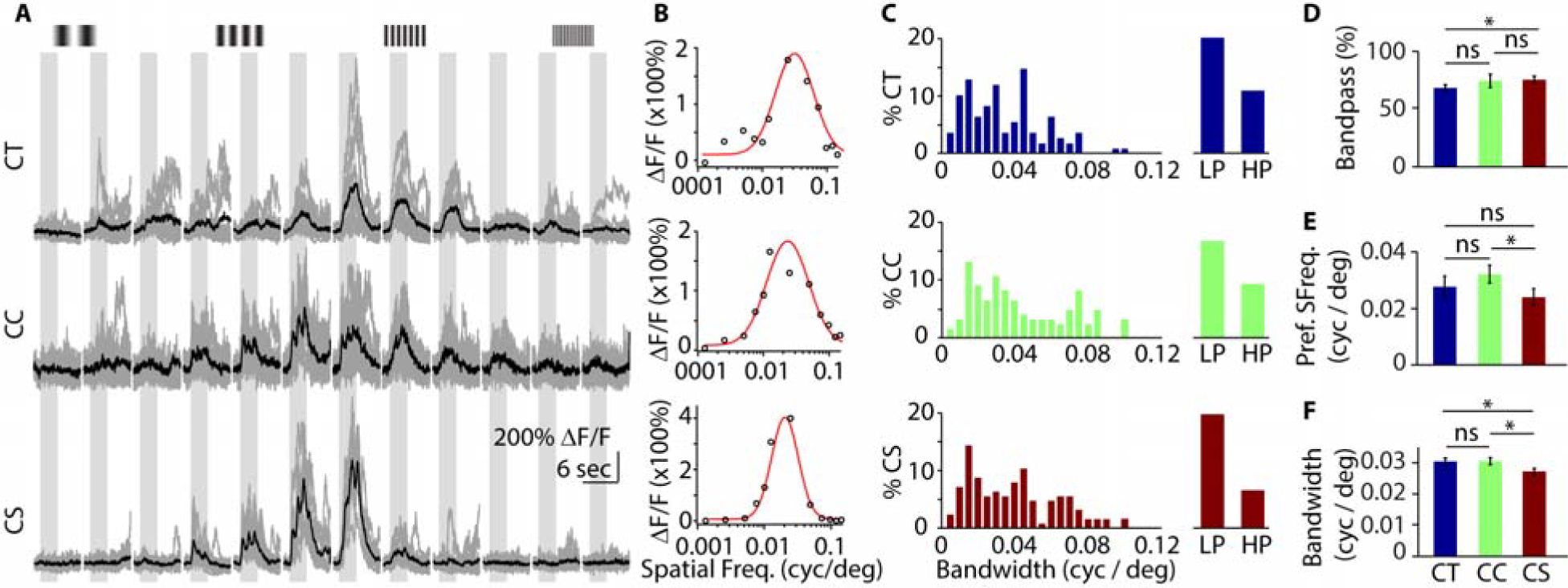
CC and CT neurons filter spatial frequencies at a broader band than CS cells. (A) Example raw (gray) and average (black) traces of a CT (top), CC (middle) and CS (bottom) neurons at varying spatial frequencies. (B) Gaussian curves (red) fit over spatial frequency data (black circles) from (A) on a log10 scale. (C) Distributions of bandwidths and fractions of low-pass (LP) and high-pass (HP) for CT, CC, and CS cells. (D) Bars represent mean ± SEM fraction of band pass cells in CT (blue) CC (green) and CS (red) populations. (E) Bars represent mean ± SEM preferred spatial frequency of CT (blue) CC (green) and CS (red) cells. (F) Bars represent mean ± SEM spatial frequency bandwidth of CT (blue) CC (green) and CS (red) cells. *: p<0.05, Student’s t-test, semi-weighted statistics (see Methods).

We next compared the orientation tuning of the three L5 subpopulations by presenting sinusoidal drifting gratings at 100% contrast in 12 different orientations. All three groups exhibited orientation selective responses (Fig. 2A-B, Fig. S4A-B), and we therefore calculated an orientation selectivity index (OSI, see Methods). Across the three populations, CT cells had a significantly lower mean OSI (0.351 ± 0.021, n=158 cells, 6 animals) than CC (0.42 ± 0.018, n=193 cells, 9 animals, p=0.0071, Student’s t-test) or CS (0.433 ± 0.011, n=169 cells, 5 animals, p=0.0003, t-test) cells, while the latter two were not significantly different (p=0.2796, t-test; Fig. 2C-D). We also calculated orientation tuning width by fitting the data with a flat top von Mises function (see Methods). Cells deemed over-fitted (extremely narrow tuning with low OSI, Fig. S4C) or yielding goodness-of-fit R^2^ values <0.4 were rejected from further analysis. Tuning widths were in good agreement with OSI measures, as CT cells had significantly broader tuning (37.675±1.796 degrees, n=123 cells, 6 animals) than either CC (32.962±1.84 degrees, n=169 cells, 9 animals, p=0.0334, Student’s t-test) or CS (33.16 5±1.4 degrees, n=152 cells, 5 animals, p=0.0263, t-test) cells, whereas CC and CT cells did not differ (p=0.4658, t-test; Fig. 2E). Similar results were found with an alternative measure of orientation tuning (Fig. S4D). As with previous findings in non-human primates (Ringach et al., 2002), we found that the OSI is a good predictor of the tuning width for individual cells (Pearson’s r=0.4118, p<0.001, Fig. S4C). Overall, these data indicate that, as a population, CT neurons are more broadly orientation tuned than either CS or CC neurons.

In a subset of experiments, we characterized the spatial frequency preferences of identified L5 PNs (Fig. 3A and Fig. S5A). Data were plotted on a log-scale and fit with a Gaussian function, allowing us to calculate the preferred spatial frequency and the bandwidth of each cell (Fig. 3B-C). Only cells with goodness of fit R^2^>0.4 were considered for further analysis. Cells were characterized as either low-pass, high-pass, or band-pass (see Methods, Fig. S5B,C). For all three L5 populations, the majority of cells were band-pass (Fig. 3C). Furthermore, we found that CC cells exhibited higher spatial frequency preference (0.032±0.003 cyc/deg, n=176 cells, 9 animals) than CS cells (0.024±0.003 cyc/deg, n=170 cells, 5 animals, p=0.0293, Student’s t-test), but were not significantly different from CT cells (0.028 ± 0.004 cyc/deg, n=168 cells, 6 animals, p=0.165, t-test). CT and CS cells did not differ (p=0.2177, t-test; Fig. 3E). Notably, spatial frequency bandwidth was significantly broader for CT cells (0.303 ± 0.011 cyc/deg, p=0.0194, t-test) and CC cells (0.303 ± 0.011 cyc/deg, p=0.0201, Student’s t-test) versus CS cells (0.272 ± 0.01 cyc/deg, t-test), whereas CT and CC cells did not differ (p = 0.49, t-test; Fig. 3F). These findings suggest that both CT and CC cells are more sensitive to broadband spatial information in comparison to CS cells.

### Noise correlations suggest functional L5 subnetworks

Studies in brain slices suggest that different populations of L5 PNs are selectively interconnected both within and across groups (Brown and Hestrin, 2009, Lefort et al., 2009). To assess the functional correlational structure of these circuits *in vivo*, we performed pair-wise noise correlation analysis between individual cells (Fig. S6). Higher correlation coefficients are thought to indicate a greater degree of either shared synaptic connectivity or common inputs (Cohen and Kohn, 2011, Schneidman et al., 2006). Within each field of view, we calculated the pair-wise noise correlation between CTB-labeled neurons (within population) and between labeled and non-identified cells (across populations) during repeated presentation of whole-field drifting gratings (Fig. 4A-E). We found that, on average, CT cells are as strongly correlated with each other (R_CT-CT_=0.042±0.04) as with the non-identified neurons around therm (R_CT-NI_=0.04±0.004, n=14 fields of view, 6 animals, p=0.3335, paired t-test). In contrast, both CC and CS cells are more strongly interconnected within their respective population than to the surrounding non-identified cells R_CC-CC_=0.046 ± 0.004, R_CC-NI_=0.024±0.004, n=14 fields of view, 9 animals, p=0.00001, paired t-test; R_CS-CS_=0.04±0.004, R_CS-NI_=0.025±0.004, n=11 fields of view, 5 animals, p=0.0011, paired t-test; Fig. 4F). These results suggest that CT cells form promiscuous local networks, whereas CC and CS cells preferentially participate in networks within their own subpopulation.

We found that activity correlation strength in all cell groups significantly decreased with increasing inter-somatic distance (Pearson’s r ranging from -0.04 to -0.15, p<0.05 in all populations; Fig. 4G). Notably for CC and CS cells, the correlation within groups was significantly higher (p<0.05 where indicated, paired t-test) than across groups for short distances, indicating that group identity is important for the connectivity of local networks. We also found that pair-wise correlations were related to the degree of co-tuning for orientation (Fig. 4H). Again, for CC and CS cells, the correlations were higher within than across groups (p<0.05 where indicated, paired t-test). Overall, our analyses suggest that CT cells are positioned to integrate visual information across large pools of L5 neurons, whereas CC and CS are preferentially interconnected within target-specific local networks.

## Discussion

In this study, we characterized the functional properties of three PN subtypes in layer 5 of mouse V1, defined by their projection targets. We showed that CT, CS, and CC cells comprise non-overlapping populations that display differences in contrast sensitivity, orientation tuning, and spatial frequency selectivity. In general, CT cells exhibit the highest contrast sensitivity and broadest tuning for orientation and spatial frequency, similar to a previous electrophysiological study of putative CT neurons (Mangini and Pearlman, 1980). Conversely, CS cells are more narrowly tuned for visual inputs, while CC cells exhibit intermediate properties. Moreover, analysis of noise correlations suggests that CT cells are widely connected to other L5 PNs, while CC and CS cells form more circumscribed networks within their own groups. These findings shed important light on the functional diversity of information processing by a cortical output layer and indicate that information streams routed to distinct downstream targets are functionally heterogeneous.

One caveat regarding our findings is that Ca2+ signaling may not accurately reflect underlying spike activity across different cell groups, potentially due to variations in GCaMP6 expression or nonlinearity of the indicator. However, using *ex vivo* imaging, we found no differences between spiking and calcium signaling for the three groups. Moreover, we found that the Ca2+ decay kinetics *in vivo* do not differ between the CT, CS, and CC cells (see Fig. S2), suggesting that all cells express similar amounts of GCaMP6 (Higley and Sabatini, 2008). Finally, previous reports have suggested visually-evoked firing rates for L5 PNs of less than 5 Hz (Hoy and Niell, 2015, Vinck et al., 2015), well within the linear regime for GCaMP6 signaling (Chen et al., 2013, Podor et al., 2015). Thus, we do not think it likely that variation in spike-Ca2+ coupling explains the observed differences in visual tuning across populations of L5 PNs.

Previous work in brain slices has demonstrated the morphological, molecular, and electrophysiological heterogeneity of L5 PNs (Hattox and Nelson, 2007, Shepherd, 2013, Larkman and Mason, 1990, Harris and Shepherd, 2015, Kasper et al., 1994). Two major cell types have been described: thin tufted corticocortical cells (also referred to as Type B or intratelencephalic) and thick tufted corticotectal cells (also called type A or pyramidal tract). Type B cells, likely corresponding to our CC and CS cells, are thought to be located primarily in layer 5A and are characterized by wider action potentials, adapting firing properties, and the expression of the transcription factor SATB2 (Shepherd, 2013). Conversely, Type A neurons, likely corresponding to our CT cells, are thought to be located in layer 5B and exhibit narrower action potentials, bursting firing patterns, and expression of CTIP2 and FEZF2 (Hattox and Nelson, 2007, Kasper et al., 1994). Notably, in the auditory cortex of the rat, intrinsic-bursting L5 PNs have broader tuning properties than regular spiking cells (Sun et al., 2013).

Work from both *in vivo* and *ex vivo* preparations has suggested the existence of synaptically coupled subnetworks within cortical microcircuits (Brown and Hestrin, 2009, Lefort et al., 2009). For example, cells that share similar visual tuning properties exhibit higher monosynaptic connection probability (Ko et al., 2011, Kohn and Smith, 2005). In addition, paired recordings of layer 5 PNs in V1 indicate high interconnectivity between CS cells, whereas CT cells are broadly connected with multiple L5 populations (Brown and Hestrin, 2009). Here, we analyzed noise correlations, which have been used to assess functional (though not necessarily anatomical) connectivity between neurons *in vivo* (Cohen and Kohn, 2011, Hofer et al., 2011, Kohn and Smith, 2005, Ecker et al., 2010, Smith and Kohn, 2008). Our results expand these previous findings to show that both CC and CS cells exhibit strong within-group correlations, suggesting preferential connectivity among like-projecting neurons. In contrast, CT cells appear to be broadly connected both within and between groups. This divergent connectivity of CT cells is further supported by their lower modulation index, suggesting that CT cells are more complex-like. Complex cells are hypothesized to arise from the summed input from upstream simple cells (Martinez and Alonso, 2003, Hubel and Wiesel, 1962), suggesting that CT cells function generally as integrators. Finally, in agreement with previous findings (Hofer et al., 2011, Ko et al., 2011, Smith and Kohn, 2008), we show that functional connectivity of all groups is significantly correlated both with similarity of orientation tuning as well as intersomatic distance. Again, for CC and CS cells, there is greater correlation within versus across group. Thus, our results indicate that projection specificity is a key additional factor in determining functional circuit interactions.

These findings indicate that subpopulations of L5 cells relay varied information about visual stimuli to different downstream targets. This conclusion is supported by recent evidence that cells in V1 that project to different ipsilateral higher-order visual areas also convey distinct spatial and temporal information (Glickfeld et al., 2013, Jarosiewicz et al., 2012, El-Shamayleh et al., 2013, Movshon and Newsome, 1996, Andermann et al., 2011). In addition, a recent study found that different genetically-defined L5 PNs exhibit tuning differences similar to those seen in our work (Kim et al., 2015). Ultimately, this organization may provide information necessary for appropriate processing by the target structures. For example, the superior colliculus is thought to play a prominent role in orienting behaviors, where fine information about spatiotemporal stimulus properties may be unnecessary (Sahibzada et al., 1986, Dean et al., 1986). This is consistent with the high contrast sensitivity and broad tuning properties of CT cells, which may function more like “detectors”. In contrast, the striatum plays a crucial role in motor planning and reward-based learning (Graybiel and Grafton, 2015). Furthermore, higher order visual areas (e.g., V2) may play key roles in decision making about visually guided behaviors (Lee et al., 2002, Prusky and Douglas, 2004, Marshel et al., 2011). Therefore, cells projecting to these areas may require higher selectivity for visual features, functioning more like “discriminators”. Future studies are needed to investigate the behavioral contributions of these heterogeneous L5 populations.

Lastly, we note that our approach to the statistical analysis of population data was based on the inherent nested design of the study. Analyses based on individual cells (rather than animals) face an increased false positive rate for detecting significant differences (Galbraith et al., 2010, Cochran, 1937). To address this issue, we used a statistical approach that compares means across animals (DerSimonian and Laird, 1986, Chung et al., 2013) (Methods), with individual means weighted by the variance within and across groups. This method is commonly used in random-effects meta-analyses and reduces the false-positive rate while maintaining statistical power within acceptable limits (Aarts et al., 2014). This is an especially important analytical tool for multiphoton data sets that typically include many tens or hundreds of cells per mouse, but do not involve large numbers of animals.

In conclusion, our findings indicate that despite physical co-mingling of cell bodies, subpopulations of V1 neurons form specific functionally interconnected networks in L5 that are capable of extracting varied feature information about the visual world and relaying this information to different downstream targets.

## Experimental procedures

### Animals

Adolescent (6-8 week) wild type C57/bl6 mice (Charles River Laboratories) were used in accordance with the Yale Institutional Animal Care and Use Committee and federal guidelines.

### *In vivo* imaging

GCaMP6s was expressed in V1 using an adenoassociated virus vector (AAV2*-hSynapsin1-*GCaMP6s, serotype 5, University of Pennsylvania Vector Core). Projection-specific subtypes of L5 PNs were labeled using CTB-Alexa Fluor-555 injected into the SC, dStr, or cV2. Imaging was performed 25-30 days after injection under light isoflurane anesthesia through an acutely implanted glass cranial window. Imaging was performed using a resonant scanner-based two-photon microscope (MOM, Sutter Instruments) coupled to a Ti:Sapphire laser (MaiTai DeepSee, Spectra Physics) tuned to 940 nm for GCaMP6 and 1000 nm for CTB-Alexa Fluor 555. Images were acquired using Scanimage 4.2 (Vidrio Technologies) at ~30 Hz from a depth of ~450-600 μm relative to the brain surface. Visual stimuli consisted of full-screen sinusoidal drifting gratings with a temporal frequency of 1 Hz and with varied contrast, orientation, and spatial frequency. For all experiments, visual stimuli were 3 seconds in duration and separated by an inter-stimulus interval of 5 seconds.

### Data analysis

Analysis was performed using custom-written routines in MATLAB (The Mathworks) and IgorPro (Wavemetrics). Regions of interest (ROIs) corresponding to single cells were selected as previously described (Chen et al., 2013). Ca2+ signals in response to visual stimuli were averaged and expressed as ∆F/F. A cell was classified as visually responsive if the Ca2+ signals during stimulus presentation were statistically different from the signals during five blank periods (p<0.05, ANOVA test) and larger than 10% ∆F/F.

The modulation index (MI) for each individual cell was determined by fitting data with a sine function and normalizing the peak-to-trough amplitude by the mean total Ca2+ response. Contrast response curves were fit by a hyperbolic ratio function (Contreras and Palmer, 2003). The orientation selectivity index (OSI) was calculated as 1 - circular variance (Ringach et al., 2002). Orientation tuning bandwidth was measured as the half width at 1/sqrt2 of a flat-top von Mises function fit to the data. For spatial frequency tuning, data were plotted on a loglO-frequency scale and fit with a Gaussian function. Cells were classified as low pass or high pass if the low or high end of the tuning curve, respectively, failed to cross the half maximum point. For all analyses that required curve fitting, cells were only included if the goodness-of-fit yielded a R^2^>0.4. Noise correlations were calculated as the partial correlation coefficient between pairs of cells.

### Statistical analysis

For most analyses, we developed a method of using semi-weighted estimators to compare individual animals, rather than cells (Chung et al., 2013, DerSimonian and Laird, 1986). This approach minimizes false positives while maintaining statistical power (Aarts et al., 2014). We used this semi-weighted estimator to calculate the statistical significance of the difference between cell populations using a standard Student’s t-test. The only exception to this was the noise correlation analysis in Figure 4, where we used the weighted estimator to reflect the pair-wise nature of the comparisons.

**Figure 4.**
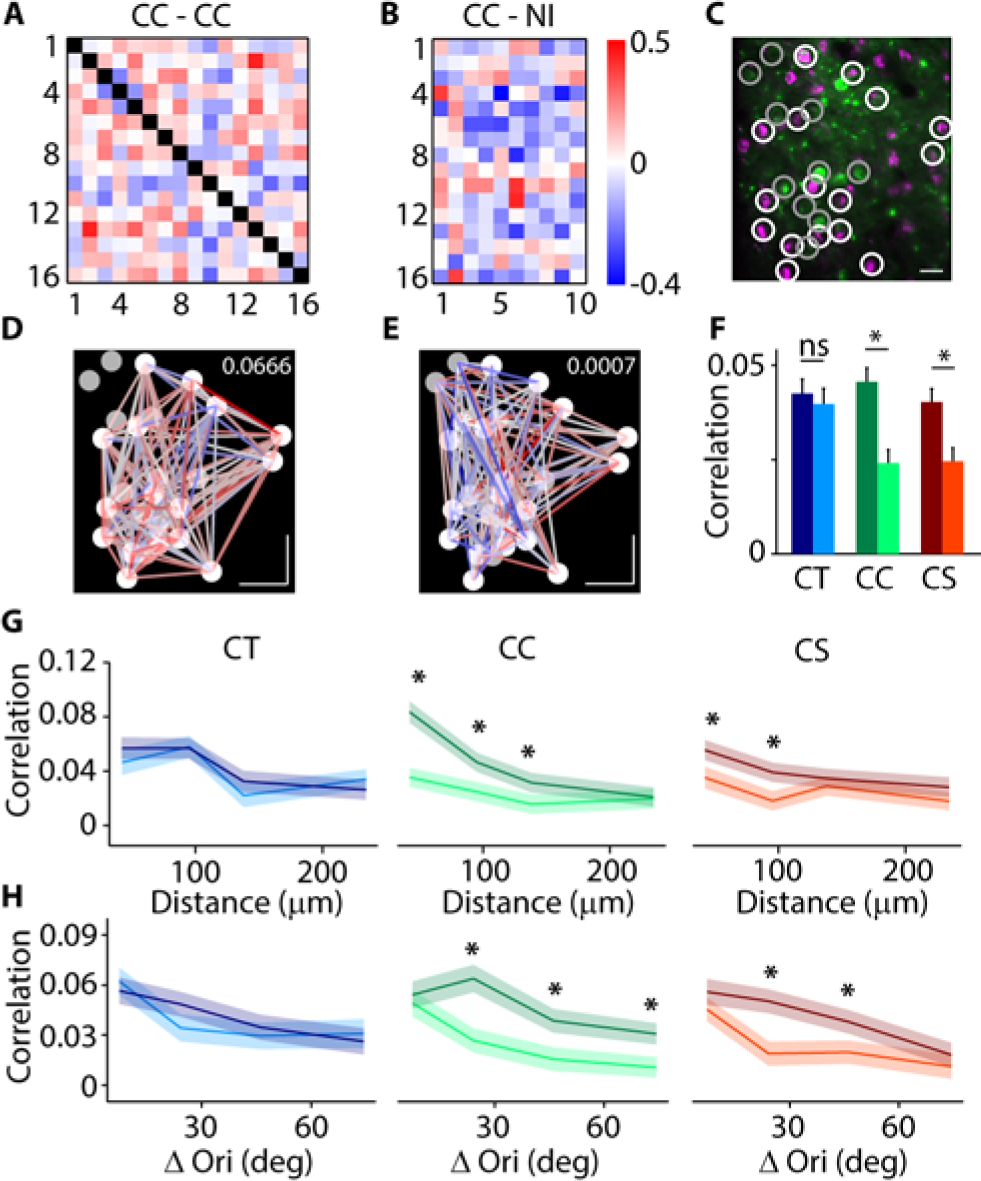
CC and CS neurons form local subnetworks. (A) Heat map showing the strength of partial noise correlations between pairs of labeled CC neurons within an example field of view. (B) Heat map showing partial noise correlations between pairs of labeled CC and non-identified (NI) neurons in the same field of view as in (A). (C) 2-photon fluorescent image of the field of view in (A) and (B) highlighting visually responsive CC (white circles) and NI (gray circles) neurons. (D) Web graph showing the connections and correlation strength between CC neurons in the same field of view as in (A-C). (E) Web graph showing the connections and correlation strength between CC and NI neurons in the same field of view as in (A-D). (F) Bars representing mean ± SEM correlation strength between CT-CT (dark blue), CT-NI (light blue), CC-CC (dark green), CC-NI (light green), CS-CS (dark red) and CS-NI (light red) cell pairs. *: p<0.05, paired t-test. (G) Change in correlation strength with distance between CT-CT (dark blue), CT-NI (light blue), CC-CC (dark green), CC-NI (light green), CS-CS (dark red) and CS-NI (light red) cell pairs. *: p<0.05, paired t-test. (H) Change in correlation strength related to the degree of co-tuning for orientation between CT-CT (dark blue), CT-NI (light blue), CC-CC (dark green), CC-NI (light green), CS-CS (dark red) and CS-NI (light red) cell pairs. *: p<0.05, paired t-test.

## Author contributions

G.L, J.A.C, and M.J.H. designed and G.L. and L.T. performed the experiments. G.L, M.A.V. and L.T. analyzed the data. G.L., J.A.C, and M.J.H. wrote the paper.

## Acknowledgements

The authors thank members of the Cardin and Higley labs for comments during the preparation of this manuscript. Special thanks to Daniel Barson for the neuropil subtraction script. The work was funded by grants from The Brain and Behavior Research Foundation (G.L., J.A.C., M.J.H.), the NIH: MH099045 (M.J.H.) and EY022951 (J.A.C.), the Alfred P. Sloan Foundation (J.A.C., M.J.H.), the Whitehall Foundation (J.A.C.), the Klingenstein Foundation (J.A.C., M.J.H.), the McKnight Foundation (J.A.C.), and a Rubicon Grant (Netherlands Organization for Science) and a Human Frontiers postdoctoral fellowship award to M.A.V.

